# Patient-Specific Pharmacogenomics unveils xCT as a Key Regulator for Druggable Metabolic and Proliferation Pathways in Colon Cancer

**DOI:** 10.1101/2025.02.11.637634

**Authors:** Marco Strecker, Keren Zohar, Martin Böttcher, Thomas Wartmann, Henry Freudenstein, Maximilian Doelling, Mihailo Andric, Wenjie Shi, Or Kakhlon, Katrin Hippe, Beatrix Jahnke, Dimitrios Mougiakakos, Franziska Baenke, Daniel Stange, Roland S Croner, Michal Linial, Ulf D Kahlert

**Affiliations:** Molecular and Experimental Surgery, Clinic for Visceral-, General, Vascular and Transplantation Surgery, University Medicine Magdeburg, Otto-von Guericke University Magdeburg, Germany; Department of Biological Chemistry, Faculty of Science, The Hebrew University of Jerusalem, Jerusalem, Israel; Clinic for Hematology and Oncology, University Medicine Magdeburg, Otto-von Guericke University Magdeburg, Germany; Department of Neurology, The Agnes Ginges Center for Human Neurogenetics, Hadassah-Hebrew University Medical Center, Jerusalem, Israel; Faculty of Medicine, The Hebrew University of Jerusalem, Jerusalem, Israel; Institute for Pathology, University Medicine Magdeburg, Otto-von Guericke University Magdeburg, Germany; Department of Visceral, Thoracic and Vascular Surgery, Medical Faculty and University Hospital Carl Gustav Carus, Technische Universität Dresden, Dresden, Germany; National Center for Tumour Diseases Dresden (NCT/UCC), a partnership between DKFZ, Faculty of Medicine and University Hospital Carl Gustav Carus, TUD Dresden University of Technology, and Helmholtz-Zentrum Dresden - Rossendorf (HZDR), Dresden, Germany

**Keywords:** xCT, patient-derived organoids, ferroptosis, chemotherapy resistance, tumor metabolism

## Abstract

Colorectal cancer (CRC) represents the third-leading cause of cancer-related deaths. Knowledge covering diverse cellular and molecular data from individual patients has become valuable for diagnosis, prognosis, and treatment selection. Here, we present an in-depth comparative mRNA-seq and microRNA-seq analysis of tissue samples from 32 CRC, pairing tumors with adjacent healthy tissues. The differential expression gene (DEG) analysis revealed an interconnection between nutrients, metabolic programs, and cell cycle pathways. We focused on the impact of overexpressed SLC7A11 (xCT) and SLC3A2 genes which compose the cystine/glutamate transporter (Xc-) system. To assess the oncogenic potency of the Xc-system in a cellular setting, we applied a knowledge-based approach for analyzing gene perturbations from CRISPR screens across various cell types as well as using a variety of functional assays in five primary patient-derived organoid cell models to functionally verify our hypothesis. We identified a previously undescribed cell surface protein signature predicting chemotherapy resistance and further highlighted the causality and potential of pharmacological blockage of ferroptosis as promising avenue for cancer therapy. Biological processes such as redox homeostasis, ion/amino acid transporters and de novo nucleotide synthesis were associated with these co-dependent genes in patient specimens. This study highlighted a number of overlooked genes as potential clinical targets for CRC and promotes stem cell-based, patient-individual *in vitro* model systems as a versatile partner platform to functionally validate in silico predictions, with focus on SLC7A11 and its associated genes in tumorigenesis.

## Introduction

Colorectal cancer (CRC) accounts for 10% of global cancer cases, making it the third most common cancer and the third leading cause of cancer-related deaths in the USA. Access to extensive cancer patient data enables evaluating of crucial predictive biomarkers needed for optimizing treatment choices. Early diagnosis is crucial for improved survival, currently based on clinical features like age, family history, tumor location, size and TNM staging (Vega *et al*, 2015). Genetic alterations, particularly recurrent mutations, are also tested for higher precision (Testa *et al*, 2018). While noninvasive tests (e.g., blood and stool tests) show promise, their lack of mechanistic or cellular interpretation limits their use (Zygulska & Pierzchalski, 2022). Surgery is the primary treatment for most CRC patients with chemotherapy being administered for advanced disease (Biller & Schrag, 2021). Although the analysis of data from large scale, open access data platforms from clinical tumor samples has accelerated our gain of knowledge on different neoplasms, they often lack proper control dataset or have limitations in the completeness of corresponding clinical metadata, especially post-surgery clinical follow-up. Nevertheless. multi-omics approaches, including diverse cellular and molecular data, has become valuable in influencing CRC diagnosis, prognosis, and treatment selection (Menyhárt & Győrffy, 2021).

Studying tumor cells’ metabolic demands, and their capacity to cope with stress, along with characterizing cancer immune microenvironment, can enhance therapy precision (Li *et al*, 2020; Li *et al*, 2021). In any living system, amino acids accessibility of amino acids is crucial for energy production, translation efficiency, and redox homeostasis (Vučetić *et al*, 2017). It has been shown that cancer cells need large amounts of cysteine and glutathione (GSH) to neutralize the increased intracellular reactive oxygen species (ROS) (Bonifácio *et al*, 2021). Cystine starvation induces cell death that can be rescued by antioxidants. Most cancer cells rely on the Xc-heterodimeric amino acid transporters system, consisting of SLC7A11 (xCT) and SLC3A2 (heavy chain 4F2hc) (Koppula *et al*, 2021). This system mediates cystine uptake in exchange for intracellular glutamate (Lin *et al*, 2020). As the extracellular environment of cells is strongly oxidizing, cysteine is rapidly oxidized to cystine (Daher *et al*, 2020). The Xc-system imports cysteines, which serve as precursors for reduced glutathione (GSH) synthesis (Lin *et al*., 2020). GSH is vital for detoxification, maintaining intracellular redox balance, reducing hydrogen peroxide and other oxygen radicals, and serving as a thiol donor to proteins. While other transporters that can partially compensate for SLC7A11 failure, it remains the major route for cystine transporter in cancer cells (Parker *et al*, 2021). In cancer stem cells (CSCs), the CD44 variant isoform (CD44v) can interact with and stabilize SLC7A11 on the cell surface (Jyotsana *et al*, 2022). While SLC7A11 was identified 40 years ago, a system view that combines expression regulation, a picture that integrates the metabolic load, redox status, and tumor microenvironment (TME) remains fragmented. Cancer cells’ nutrient dependency generally increases SLC7A11 function. High expression of SLC7A11 reduces oxidative stress in some oncogenic *KRAS*-mutant cancers, supporting cancer progression (Koppula *et al*., 2021). In many cancer types xCT supports tumor growth by preventing ferroptosis and allowing cells to cope with chemotherapy and radiation-induced oxidative stress. A correlation between high level of SLC7A11 activation and low survival rate is reported for pancreatic ductal adenocarcinoma (PDAC) and lung adenocarcinoma (LUAD) (Lin *et al*., 2020). Detection of xCT expression in colorectal cancer has also been previously reported to be associated with faster disease recurrence (Sugano *et al*, 2015), along with the correlation with immune cell infiltration into the tumor and poorer survival of patients (He *et al*, 2021). Experimentally, knockdown of *SLC7A11* led to increasing intracellular ROS, which potentially arrest tumor invasion (Cheng *et al*, 2022).

In this study, we focus on the transcriptional profiles (mRNAs and miRNAs) from 32 colon cancer patients, each analyzed by comparing their tumor to the healthy tissue. We identified strongly upregulated coding gene sets that signify mitotic cell signature from colon, cell cycle G2/M checkpoints and an additional network of dysregulated transporters leading to a metabolic burden. We focused on SLC7A11 and its functional network as an integrator of colon cancer progression. Using functional CRISPR cellular fitness analysis and survival data from large resources, we identified genes carrying clinically relevant properties. A comprehensive bioinformatic analysis explored the potential of Xc-system status for clinical decisions. While therapy responsiveness and composition of immune cells did not provide a valid signal for prediction, genes involved in nutrient supply, mitochondria and redox state were strongly indicative. Further, functional investigations using primary patient-derived organoids (PDOs) from patients with CRC showed that therapeutic responses to both standard regimens and xCT inhibitors are significantly modulated by xCT expression levels and the newly identified surface marker profile predicting for therapy response. We illustrate the importance of a multilayer analysis in exposing overlooked cellular processes and targets for improving the clinical and therapeutic management of CRC. Our project highlights the usability of PDO platform for guiding stratification of therapy resistance levels of tumors as well as supporting early-stage drug development.

## Methods

### RNA-seq analyses of 32 CRC patients

The mRNA expression levels of all genes (coding and non-coding) were determined by pairwise analysis of cancerous and healthy tissues obtained from the same patient. Total of 32 patients were analyzed with 73 deep sequencing results. Altogether, there were 36 samples marked as tumor (T) and 37 samples marked as healthy (H). Each participant provided at least one sample for T and H. For four participants the number of samples was higher. Each sample was studied by partition total RNA by size to support mRNA and miRNA transcriptomics.

Ethical approval to conduct this study was granted by the ethics committee of the medical faculty of Magdeburg (33/01, amendment 43/14).

### Analyses of CRC patients from public resources

The Limma R package (Ver 4.2.0) was used for differential expression analysis with adjusted p-value of 1e-20 (for pair-wise analysis) as significance threshold. We have applied GEPIA2 database that covers the data from The Cancer Genome Atlas (TCGA) and Genotype-Tissue Expression (GTEx) (Consortium, 2013). Box plot, violin plot, and scatter plot for selected differentially expressed genes (DEGs) were drawn by the TCGA and GTEx visualization website GEPIA2 (Tang *et al*, 2019).

### Colon cell type

Analysis of bulk RNA-seq datasets from 15 human organs including colon produced a cell type enrichment prediction atlas for all coding genes. The initial data is extracted from GTEx. The identity profiles across tissue types revealed 12 types of cells in colon by the Human Proteome Atlas (HPA; (Thul & Lindskog, 2018)). The 12 colon cells cover 1918 genes. We performed the analysis for 7 main colon cell types: Colon enterocytes (369 genes), Colon enteroendocrine cells (338 genes), Enteric glia cells (240 genes), Mitotic cells in Colon)85 genes), Endothelial cells (219 genes), Smooth muscle cells (166 genes), Fibroblasts (42 genes). Additionally, 5 immunological cell types from colon are: Macrophages (143), Neutrophils (65), Mast cells (29), T-cells (108) and Plasma cells (114).

### Bioinformatics tools and statistics

#### Statistically significance

Paired statistics for 2-group analysis was based on 2-tailed t-test. Statistical significance was also computed using the non-parametric Mann-Whitney U-test. Kruskal-Wallis tests used in single-variable comparisons with more than 2 groups. Differences with *p* <0.05 were regarded as statistically significant (unless mentioned otherwise). False Discovery Rate (FDR) was computed using the Benjamini-Hochberg method. Hypergeometric distribution test was used to obtain p-values for overlapping gene sets.

#### Gene expression density plot

Conducted using RNA-seq data from TCGA, combined with the Therapeutically Applicable Research to Generate Effective Treatments (TARGET), and the GTEx repositories using TNMplot (Bartha & Gyorffy, 2021).

#### Enrichment tests

The database of gene sets from the Molecular Signatures Database (MsigDB). A collection of hallmark gene sets is a set of 50 main processes in cells with expert curation with about 200 genes included in each hallmark set (Liberzon *et al*, 2015). Testing overrepresentation analysis by slice representation. The different colored slices indicating the hallmarks (total of 10) that are significant (using the adjusted p <0.05 as a threshold). The analysis used 6763 genes that are associated with any of the hallmarks as a reference set.

Enrichment for miRNAs was based on miRinGO, that address indirect gene targets genes through transcription factors (TFs) according to miRNA expression in specific tissues (Sayed & Park, 2023). Database of dbDEMC 3.0 (database of differentially expressed miRNAs in human cancers) covers 40 cancer types with large scale compilation of miRNA gene expression from experiments (Xu *et al*, 2022).

### Interacting networks

Physical protein-protein interaction (PPI) set is based on IntAct with over 1M interactions datapoints. High confidence interactions were based on the molecular interaction (MI) score. The MI score calculates on independent PPI evidence and complementary experimental methods. Spoke expansion refer to all pairs of bait identified interactions (Del Toro *et al*, 2022). STRING network was used based on PPI confidence score (>0.6, or as mentioned). Connectivity network excluded neighborhood, gene fusion and co-occurrence as evidence (Szklarczyk *et al*, 2021). Network of miRNA-genes TF regulation by miRNAs was extracted from EMT-Regulome (Zhao *et al*, 2017).

### CRISPR-Cas9 cell line screening

We used the pre-calculated correlation of dependency from DepMap using CRISPR/Cas9. CRISPR-Cas9 screenings are reported for 19,144 genes across 1206 cell lines (primary and established) and providing knockout fitness scores (14 days after transfection) (Dempster *et al*, 2019). Dependencies enriched in COAD were precalculated for ∼1800 genes (identified by DepMap CRISPR-Cas9 project using the Public 23Q4+Score, Chronos resource), with an expanded collection of cell lines and cancer types from BioGRID ORCS (Ver. 1.0.4) CRISPR screen (Choi *et al*, 2021). We search for genes with correlated knockout fitness (called ‘co-dependent’). A loss of fitness and a negative log fold change in the average representation of the relevant targeted sequence relative to plasmid are indicative for the gene being essential.

### Predictive analysis by gene expression level

KM Plot and ROC Plotter (Fekete & Győrffy, 2019) were used to identify gene expression-based predictive biomarkers for CRC that compiled publicly available datasets. By integrating gene expression data (RNA-seq and Chip-Seq) with chemotherapy, most genes were tested. A link of gene expression and therapy response using transcriptome-level CRC data generates a ROC plot with detailed statistics on relevance of any gene to therapy and clinical response (Fekete & Gyorffy, 2023). To identify genes with altered expression in TCGA COAD samples in view of disruptive mutations in a gene set (5 genes) we applied the GENOTYPE module of MuTarget (Nagy & Gyorffy, 2021).

### Establishment of COAD and healthy colon patient-derived organoids

Detailed reagent information is provided in **Supplementary Text S1** Freshly resected tissue samples were transported on ice to the Institute of Pathology, where a pathologist confirmed tumor presence (for CRC samples) and processed healthy tissues. For healthy colon organoids, the mucosa was separated from the submucosa and muscularis, cut into 1–2 mm fragments, and washed three times in wash buffer (DPBS with 1% antibiotic-antimycotic solution and 0.1% gentamicin sulfate). Crypts were liberated via incubation in DPBS containing 2 mM Na-EDTA for 1–2 h, followed by gentle pipetting and repeated collection of the crypt-containing supernatant. Crypt solutions were then centrifuged (200 × g, 4 °C, 5 min) and washed twice.

To establish organoids from CRC tissues, small fragments were washed three times in the same wash buffer before enzymatic digestion with 5 mg/ml Collagenase IV, 250 µl Dispase, and 1750 µl Basic Media (Advanced DMEM/F12, 1% Hepes, 1% Penicillin-Streptomycin, 1% Glutamax) in a shaking water bath at 37 °C for 15–30 min. Digestion was halted upon observing single cells and clusters, and the suspension was passed through a 100 µm cell strainer, centrifuged (300 × g, 4 °C, 5 min), and washed three times in wash buffer.

Isolated crypts or tumor cells were resuspended in Cultrex Basement Membrane Extract (BME) and seeded in 10 µl drops on pre-warmed plates, which were incubated upside down at 37 °C for 30 min to allow polymerization. Organoids were maintained in Colon Expansion Media (Basic Media supplemented with 10% R-spondin-CM, 5% Noggin-CM, 2% B27, 10 mM nicotinamide, 1.25 mM N-acetylcysteine, 10 nM gastrin, 50 ng/ml mEGF-Rec, 500 nM A-83-01, 10 µM SB202190, 100 µg/ml Primocin, and 1 µM PGE2). For healthy colon organoids, NGS-Wnt recombinant protein (final 1 nM) was added (Miao *et al*, 2020). Immediately after isolation and splitting, 10 µM Y-27632 (Rho kinase inhibitor) was included in the expansion media. The study was approved by the Ethics Committee of the Medical Faculty, University of Magdeburg (#46/22).

### PDO quality control by DNA analysis

Organoids were harvested washed with DBPS and cell pellets were frozen in liquid nitrogen and subsequently stored on -80°C until further use. Organoids were process with the QIAamp DNA Mini Kit (Quiagen # 51306) following the manufacturers protocol.

Targeted parallel sequencing was carried out using a Nextera Rapid Capture Custom Enrichment panel (Illumina, San Diego, CA, USA) covering all coding exons and flanking intronic sequence of ±20 nucleotides of target genes according to manufacturer’s instructions. Cluster generation and sequencing was implemented on an Illumina MiSeq System (Illumina). Reads were aligned and mapped to the human assembly hg19 (GRCh37). We used the varvis genomics software v.1.25.0 (Limbus Medical Technologies GmbH, Rostock, Germany) for SNV- and CNV-analysis. A coverage depth of >100x has been reached.

### Flow cytometric assessed surface expression and linked PDO metabolism

The importance of PDO for testing cellular physiology and drug screening was established (e.g., (Narasimhan *et al*, 2020)). Single cell suspensions of PDOs (n=5 from colon carcinoma, n=2 from healthy donors) were prepared. Approximately 50,000 cells per sample were subjected to various staining protocols.

For the analysis of surface markers, cells were first incubated with a viability dye (Ghost Dye™ Violet 510, Cytek) following the manufacturer’s recommendations. They were then stained with the following fluorochrome conjugated antibodies for 30 min at 4° C. PE-Cy7 rat anti-human CD44 (clone QA19A43, Biolegend), FITC mouse anti-human CD71 (clone M-A712, BD Biosciences), and CF647 rabbit anti-human SLC7A11/xCT (polyclonal, biorbyt).

Cystine uptake was assessed using BioTracker Cystine-FITC Live Cell Dye (Merck) at a final concentration of 0.5 µM. Cells were incubated for 30 min at 37 °C in 5% CO_2_ using serum-free AIM-V medium (Thermo Fisher Scientific).

The labile iron pool (Fe2+) was measured using Phen Green SK diacetate (Cayman chemicals) at a final concentration of 10 nM. Cells were stained for 15 min at 37 °C in 5% CO_2_. Phen Green fluoresces in absence of Fe2+, but its fluorescence is quenched by Fe2+, enabling semi-quantitative analysis of iron levels.

All data was recorded on a NorthernLights spectral flow cytometer (Cytek) after proper titration. Data was analyzed and visualized using FlowJo V10 (BD Bioscience).

### Analysis of drug effect to PDOs by luminescence-based quantification of cell mass viability

Organoids were expanded to a suitable size and underwent at least one passage post-thaw before drug testing, typically between passages 8 and 15. For seeding single cells, organoids were harvested and washed twice in wash buffer (centrifugation at 250 × g) to remove debris, followed by a wash in DPBS (without Mg^2+^ and Ca^2+)^. The pellet was then digested in TrypLE Express and incubated for 5 min in a prewarmed 37 °C water bath. After incubation, organoids were mechanically disrupted by pipetting, and dissociation into single cells was verified microscopically. If necessary, digestion was extended by an additional 5 min. The resulting single cells were filtered through a 40 µm cell strainer, counted, and resuspended in BME.

BME drops of 5 µl containing 2,000 cells were seeded onto 96-well plates and polymerized for 30 min at 37 °C in 0.5% CO_2_. Following polymerization, Growth medium containing a Rho kinase inhibitor (Y-27632) was added, and cells were cultured for 3 days to allow organoid formation. On Day 3, drug solutions were prepared at the desired concentrations and applied for 72 h. Viability was measured using a plate reader with luminescence detection (parameters: Attenuation OD1, no lid, appropriate integration time). After 72 h of treatment, Cell Titer-Glo 3D reagent was added, and viability was assessed following the manufacturer’s instructions.

## Results

### Pairwise analysis of samples from colon cancer patients

We have analyzed 32 colon cancer patients with 73 datasets. Each patient contributed at least two samples, one from the tumor and another from the neighboring tissue classified as tumor cell free tissue. All samples were subjected to deep sequencing for mRNA profiling (see Methods).

**Fig. 1A** shows the unsupervised partitions of all 73 samples labeled tumor and healthy (T and H, respectively). The dendrogram shows a clear partition of all samples into two main branches. The tumor (T) branch is 100% consistent (purple, 31 samples), and the second major branch is mostly composed of healthy samples (88% of 42 samples, orange), with only 5 T-labeled samples clustered with the H-samples. **Fig. 1B** shows that dimensional reduction by principal component analysis (PCA) supports a successful unsupervised partition of T and H samples with 37% of the total variance explained by PC1 and PC2. The PCA used the top 1000 ranked differentially expressed genes (DEGs). A similar successful partition of T and H was achieved by PCA, whose input consisted of the entire RNA-seq profiles (total of 13,682 identified transcripts).

**Figure 1.**
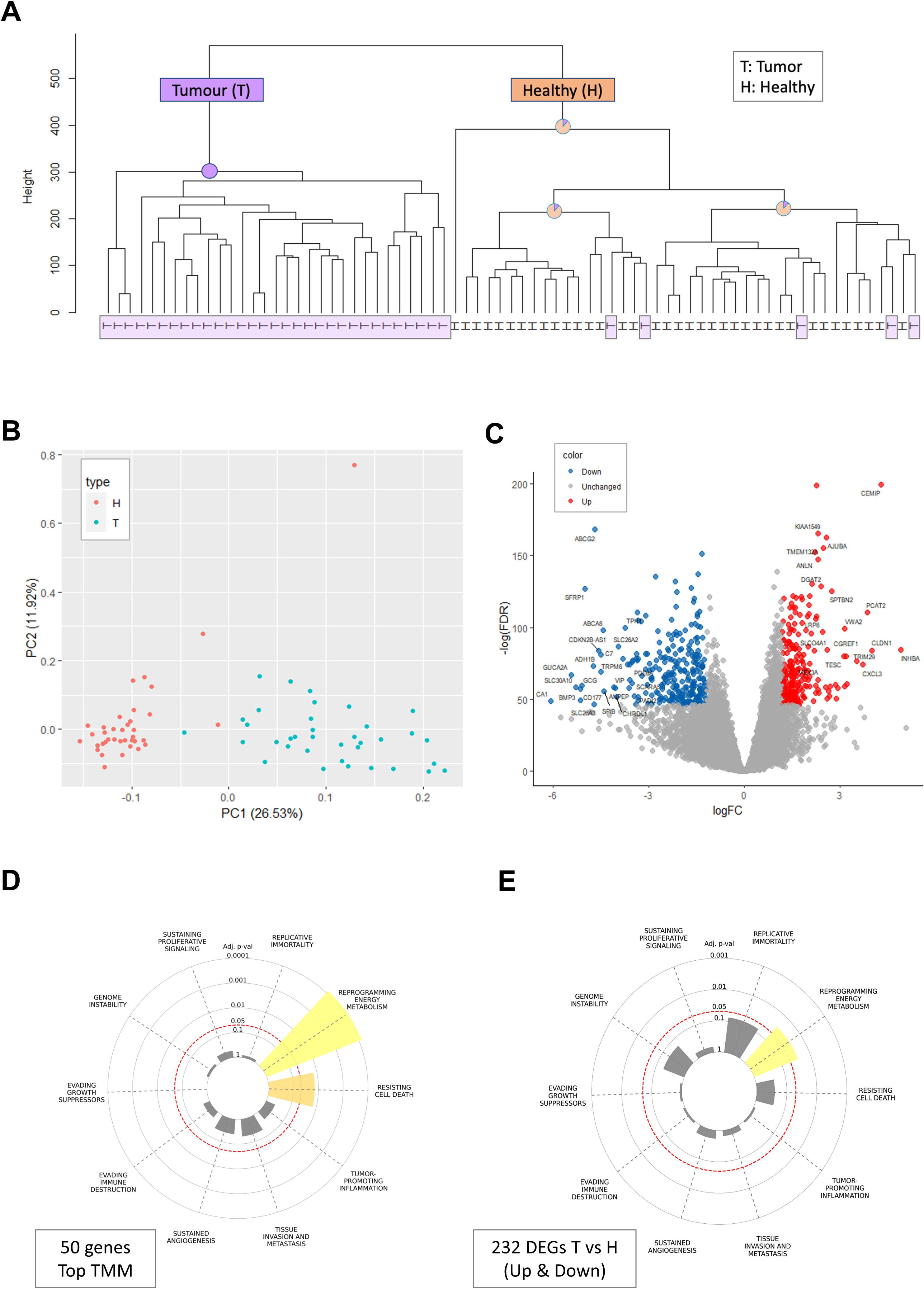
Analysis of the mRNA profiles from 32 colon cancer patients. **(A)** Unsupervised dendrogram of 73 samples from 32 participants. The main nodes are indicated by their purity for tumor (T) and healthy (H) colored purple and orange, respectively. The T samples of the dendrogram tree are highlighted with light purple background. **(B)** PCA for 73 samples, based on the top 1000 differentially expressed genes (DEGs) colored by T and H with red and blue, respectively. The variance explained are indicated for PC1 and PC2. **(C)** Volcano plot representation of DEG analysis from RNA-seq of T versus H for the samples described in A. Red and blue points mark the genes with significantly increased or decreased expression in T relative to H, respectively. Representative significant genes are indicated. **(D)** Top 50 expressing genes (RNA-seq, normalized by trimmed mean of the M-values (TMM) tested for enrichment for any of the 10 cancer hallmarks. **(E)** DEGs (up and down; 323 genes). The significantly enriched hallmarks are colored.

Next, we analyzed the consensus DEGs from all patients. Each patient was analyzed with respect to their own health-labelled sample. Single samples from the tumor and healthy tissue were normalized and compared internally (according to the number of samples available). Altogether, we performed global analyses of 32 pre-analysis patients to confirm high statistical significance and a minimal fold change threshold per gene. Specifically, the analysis was restricted to genes with a minimal statistics of FDR p-value <1e-20, with a minimal average expression of 10 counts per million (CPM) and limited the analysis for coding genes (*i.e.,* 92% of all mapped transcripts). Such filtration reduced the 13,682 unique gene transcripts to 9,045 genes that were further analyzed (**Fig. 1C**, Supplementary **Table S1)**.

We tested the results of the RNA-seq analysis to identify a signature for any of the 10 known cancer hallmarks (Hanahan, 2022). The highly expressed genes already identified significant hallmarks such as ‘reprogramming energy metabolism’ (p-value <1e-06) and ‘resisting cell death’ (adjusted p-value 1.4e-02; **Fig. 1D**).

For clinical relevance, it is essential to focus on consistent expression difference in T to H samples. To this end, we reanalyzed DEG at a relaxed threshold. For enrichment of cancer hallmarks DEG were selected with FDR ≤1e-20, a minimal fold change of 2.3 (*i.e.,* log(FC) ≥|1.2|), and a minimal average expression of 10 CPM (Supplementary **Table S2**). The signature for ‘reprogramming energy metabolism’ remained significant (adjusted p-value 2.7e-02) (**Fig. 1E**). We concluded that a signature of metabolic programming dominated DEG from the CRC patients.

### Colon cancer DEGs reveal hallmarks of cell cycle and metabolic program in cancer samples

Major cellular biological processes (see Methods) cover the preselected 50 gene sets (MSigDB). **Table 1** lists the most enriched hallmark sets (Adjusted p-value <1e-05). Several observations can be made based on the results in **Table 1**. Firstly, the stronger enrichment is for a set of genes encoding cell-cycle related targets of E2F transcription factor and G2/M checkpoint which are exclusively composed of upregulated genes. Moreover, for most significantly enriched hallmarks, the associated genes were consistent by their expression trend (i.e., either up- or downregulated). An exception is a hallmark called ‘estrogen response-late’ that shows a mixture of up- and downregulated genes. Lastly, the metabolic hallmarks are primarily associated with downregulated genes. For a list of enriched hallmarks along with their associated genes (total 27 sets) see Supplementary **Table S3**.

**Table 1.** Enrichment analysis of DEGs for the set of MSigDB hallmarks

| Hallmarks (H) gene set (MsigDB) | N in geneset | # Up | # Down | Adj. p-val |
| --- | --- | --- | --- | --- |
| H: E2F_TARGETS | 194 | 30 | 0 | 6.93E-18 |
| H: G2M_CHECKPOINT | 192 | 28 | 0 | 2.87E-16 |
| H: MTORC1_SIGNALING | 193 | 15 | 2 | 8.65E-07 |
| H: XENOBIOTIC_METABOLISM | 196 | 4 | 13 | 8.65E-07 |
| H: ESTROGEN_RESPONSE_LATE | 196 | 9 | 8 | 8.65E-07 |
| H: MITOTIC_SPINDLE | 197 | 15 | 2 | 8.65E-07 |
| H: FATTY_ACID_METABOLISM | 155 | 2 | 13 | 9.38E-07 |
| H: KRAS_SIGNALING_UP | 195 | 11 | 4 | 1.56E-05 |

The results from **Table 1** can be broadly classified into two larger themes: cell cycle-related (*e.g.,* mitotic spindle, G2M checkpoint and genes encoding cell cycle E2F) and nutrients and metabolic programs (*e.g.,* fatty acids synthesizing, genes involved in processing drugs and other xenobiotics). **Fig. 2A** tests gene overlap of the hallmark sets that belong to these main themes. With 20 genes overlapping hallmark set of cell cycle, 8 genes signifying mTOR signaling (including SLC7A11) and 4 genes overlap both sets.

**Figure 2.**
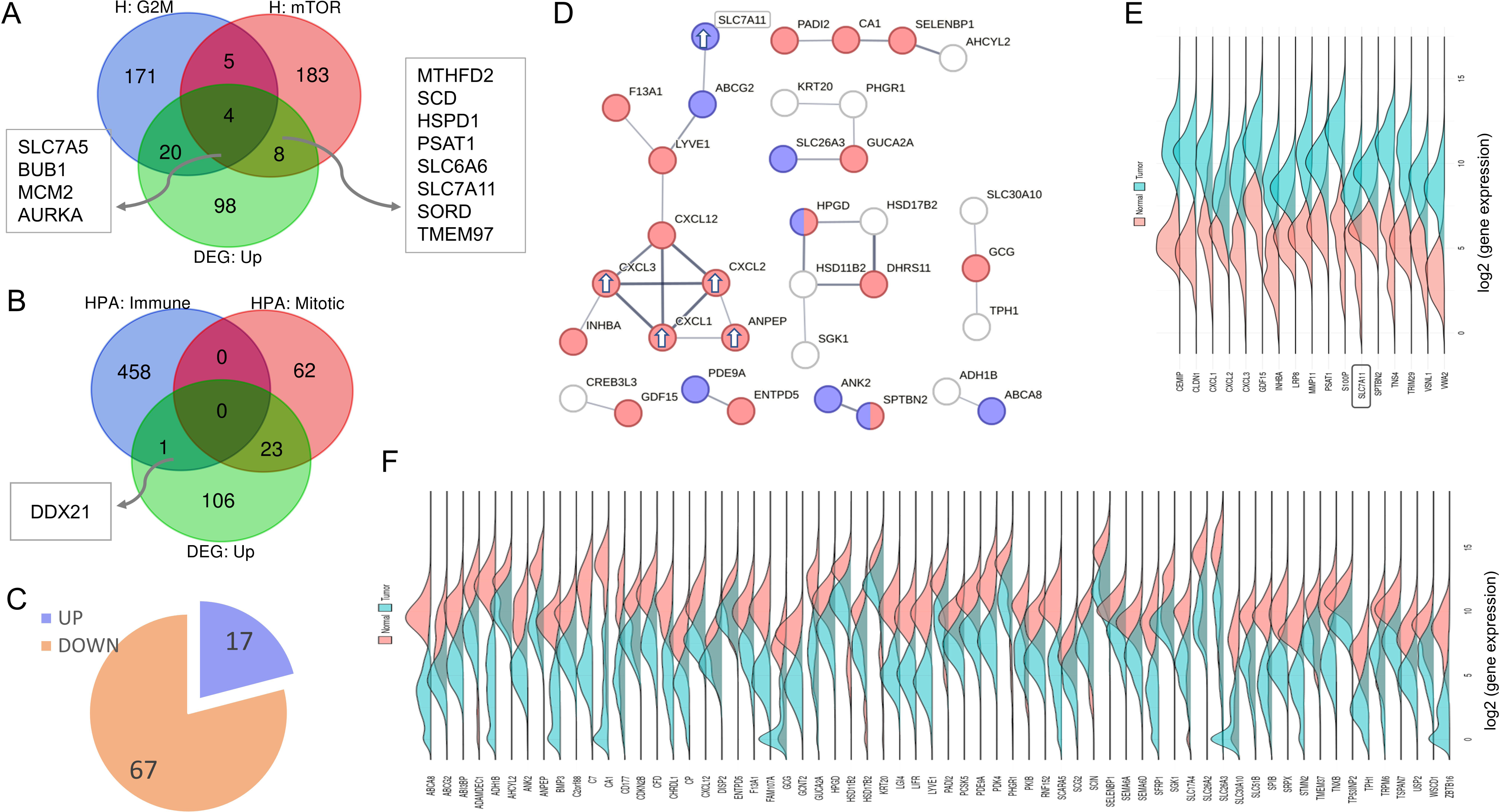
Upregulated DEGs and cellular processes. **(A)** Venn diagram of the upregulated DEGs (130 genes) and the cancer hallmark sets of ‘cell cycle G2M checkpoint’ and ‘mTORC1 signaling’. The overlap genes are listed. **(B)** Venn diagram of the upregulated DEGs (130 genes) and the colon cell types (see Methods). **(C)** Pie chart of the top 84 DEGs filtered by threshold of FC ≥|5|, and partitioned to up and downregulated genes. **(D)** STRING based network (confidence threshold 0.5). Colored are genes that are annotated by GO cellular component annotation (GO_CC) as extracellular and plasma membrane regions (red and blue, respectively). The upregulated genes in the largest connected component are indicated by white arrows. The SLC7A11 gene is highlighted. (**E)** Density plot analysis of the 17 upregulated DEG (alphabetic order). Expression density plots of healthy and tumor samples are in pink and blue, respectively. The SLC7A11 gene is marked. **(F)** Density plot analysis of the 67 downregulated DEGs (in alphabetic order).

### The upregulated genes strongly identified subpopulation of mitotic cell types

The colon is a complex tissue composed of numerous cell types. We tested the set of most significant upregulated genes (total 130, Supplementary **Table S4**) with respect to the 12 characterized cell types that are signified by enriched gene sets, based on Human Proteome Atlas (HPA) classification (see Methods). A significant overlap was detected only to mitotic cells, with 23 DEGs overlapping 85 mitotic cell enriched genes (enrichment p-value 3.5e-05). The 23 genes that are shared by the upregulated DEGs and the mitotic cells **(Fig. 2B)** produces highly connected protein-protein interaction (PPI) network (STRING, confidence p-value <1.0e-16). Among these genes are kinesin-like proteins that act in chromatid segregation (*KIF2C*, *KIF14*, *KIF20A*), genes that participate in cell cycle via DNA repair mechanism (*RAD51AP1*, *EXO1*, *BRCA2*), and checkpoint control genes that act in DNA replication (*TPX2, BUB1*, *NUF2*, *CDC6*).

Notably, among the colon-centric immune unified cell types (total 459 genes, see Methods), only *DDX21* (DEG p-value FDR 5.30e-46) was detected (**Fig. 2B)**. In colon cancer, the knockdown of *DDX21* inhibited cell growth by activating CDK1. DDX21 was postulated to mediate this effect via chromatin modulation of the CDK1 promoter (Lu *et al*, 2022). The CDK1 was also identified among the upregulated genes overlapping with the mitotic signature. Colon cell types (1918 genes, 12 cell types see Methods) are listed in Supplementary **Table S4**.

### Enrichment of extracellular and plasma membrane among the strongest DEGs

We then tested the possibility of identifying tumor versus healthy genes and focused on the subset of genes showing the most extreme differential expression signals. To this end, we analyzed a subset of 84 DEGs with fold change (FC) of >|5|. While this is an arbitrary threshold, it captures the maximally responding DEG group. **Fig. 2C** shows that at this threshold, 80% of the genes were downregulated and only 20% were upregulated.

**Fig. 2D** shows a connectivity map of these DEGs (STRING confidence score ≥0.5; minimal ≥2 connected genes). The largest subnetwork (10 nodes) with SLC7A11 transporter, genes that function in mitochondria and a set of secreted cytokines included upregulated genes (marked with arrows). The other subgraphs include the downregulated genes (**Fig. 2D**). Most connected genes (70%) were assigned with either extracellular regions (GO cellular component, p-value =3e-06; colored red) or plasma membrane region (p-value 0.003; colored blue). Notably, there were 6 transporters among the 84 upregulated genes, but *SLC7A11* was strongly upregulated while the rest of the transporter encoding genes were downregulated (Supplementary **Table S2**). These findings highlight the importance of plasma membrane signaling and extracellular communication.

To validate that the DEGs from our study are in agreement with the large-scale available data from cancer resources, we performed density plot analysis for the upregulated **(Fig. 2E**) and downregulated (**Fig. 2F**) genes with data extracted from TCGA. The results show a complete agreement in the list of all 84 DEGs (Supplementary **Table S2**) regarding the expression level trends in healthy and tumor samples of our CRC cohort.

### SLC7A11 and its interactors exhibit a coordinated upregulated expression in CRC

The enrichment of metabolic hallmarks (**Table 1**), and the results of **Fig. 2** indicating the plasma membrane enrichment, led us to focus of the Xc- system, consisting of *SLC7A11* (xCT) and *SLC3A2* (heavy chain 4F2hc). This transport system governs the intracellular redox balance in cancer cells. **Fig. 3A** lists validated direct partners of SLC7A11 (see Methods), along with the expression levels of these direct interacting partners in healthy (H) and tumor (T) samples for all 32 CRC patients **(Fig. 3B)**. The KRTAP1-1 and KRTAP1-3 were below the expression threshold in CRC samples. The IFT70B (also called TTC30B) was poorly expressed in CRC samples and showed no difference between expression in T and H. These genes were not further analyzed. SLC7A11, SLC3A2 which composes the Xc- system, and CD44, a stem cell marker, that stabilizes SLC7A11 on cell surface were significantly increased in T vs H. To generalize and validate our findings, we compared expression trend of SLC7A11 among publicly available COAD (461) and READ (172) samples from TCGA and confirmed the significant overexpression of SLC7A11 transcripts in in T vs H for **(Fig. 3C).**

**Figure 3.**
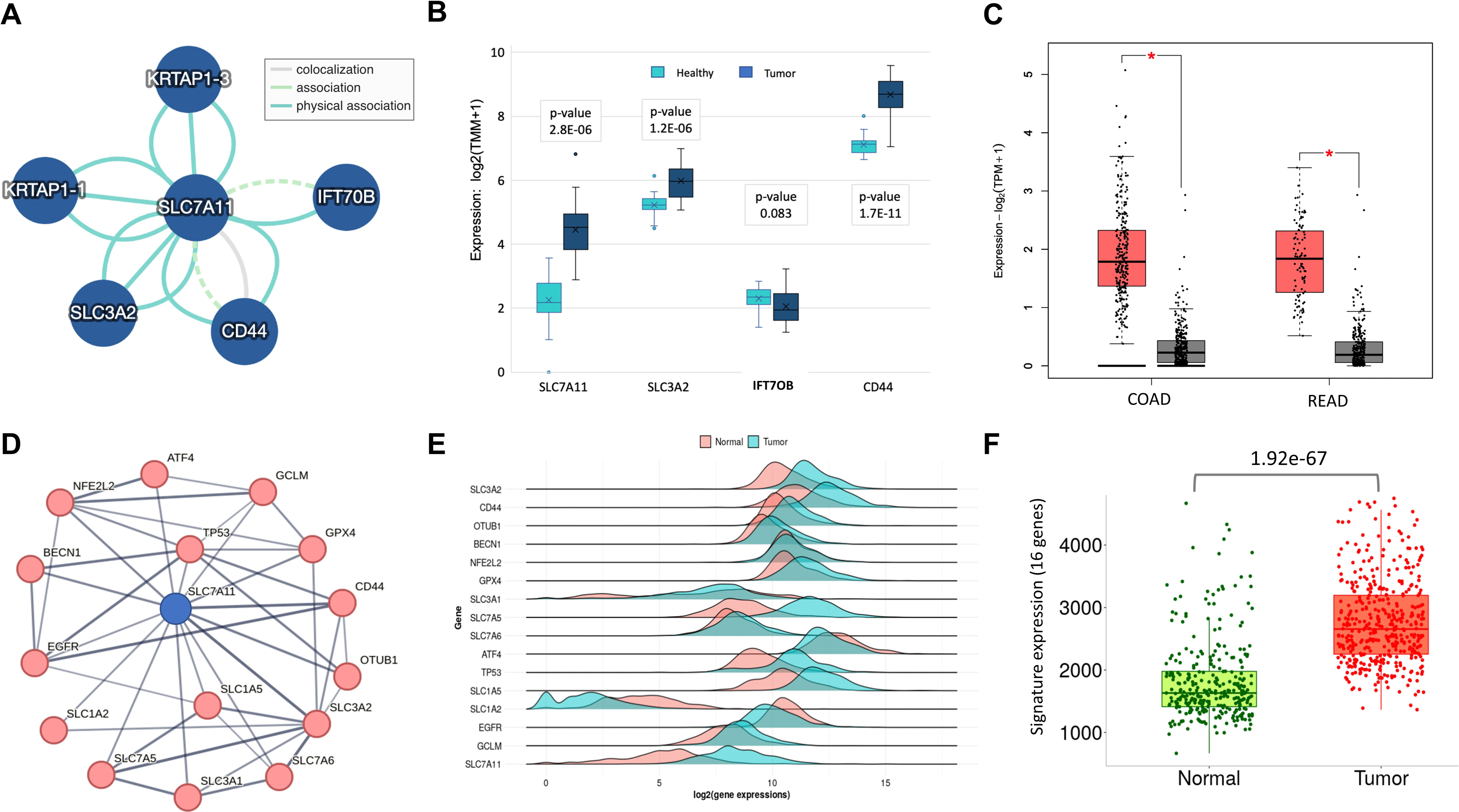
Signature of SLC7A11 network in colon cancer. **(A)** Direct physical associations of SLC7A11 (IntAct with MI score >0.5). Each partner was identified by interaction multiple methods. Dashed edge marks spoke expansion (see Methods). **(B)** Box plot of the 32 CRC patients with the expression of a subset of genes from (A). (**C)** Box plot of log expression measured by transcripts per million (TPM) for SLC7A11 from TCGA for COAD (461) and rectum adenocarcinoma (READ, 172) samples. Tumor and healthy are colored red and gray, respectively. (**D)** Interacting genes centered by SLC7A11 according to STRING (confidence score >0.6), limited to the most reliable PPI connectivity (total 16 genes). **(E)** Density plots for 16-core genes from (A). Healthy and tumor samples are marked in pink and blue, respectively. (**F)** Box plot for the signature of all SLC7A11 16-core gene set for healthy (normal, green) and tumor (red). Each dot represents a datapoint from COAD samples.

The set of genes that were strongly connected to SLC7A11 across multiple tissues were further analyzed (STRING confidence score ≥0.7; coined SLC7A11-core set, 16 genes). SLC7A11 PPI display PPI association with SLC3A2, CD44 (direct, as in **Fig. 3A**) but also with additional transporters (6) from the SLC family. Notably, the expression profiles of SLC genes with regard to the metabolic remands in cancer tissues suggest that their link to tumor immune microenvironment (TME) and drug response (Lavoro *et al*, 2023). Among the SLC7A11-core set there are other metabolic-related genes. For example, OTUB1, a specific deubiquitylating enzyme with a cysteine protease activity, BECN1 (Beclin 1) that regulates vesicle-trafficking processes, autophagy, and apoptosis. Other core genes (ATF4, NFE2L2 and GTX4) act under starvation, oxidation and ER stress. Specifically, the transcription factor (TF) ATF4 acts to induce various amino acid transporters and enzymes that determine the metabolic state of cells (e.g., redox balance, energy production, nucleotide synthesis). Other core genes are directly associated with cancer progression (TP53 and EGFR) that drive cell migration, differentiation and cell growth (**Fig. 3D**).

**Fig. 3E** shows a density plot for all 16 core-SLC7A11 genes according to COAD expression levels from TCGA. The expression levels of most core genes are higher in tumor relative to healthy samples. However, SLC1A2, EGFR and ATF4 display an opposite expression trend. The analysis for all 16-SLC7A11 core genes among our 32 CRC cohort is shown in Supplementary **Fig. S1.** We have confirmed that the overall signature of all 16 core-SLC7A11 genes remain highly significant for the difference of tumor relative to healthy tissue for the entire TCGA cohort of COAD (Mann-Whitney p-value 1.9e-67, **Fig. 3F**). We concluded that SLC7A11-core set is a hub for processes of mitotic cells and metabolic balance in among our CRC patients and this signal is validated across 630 samples from TCGA (COAD and READ).

### Knowledge-based inspection of SLC7A11 determines its oncogenic potency

We sought to identify functional network of SLCA11 correlated genes by considering gene perturbations in a cellular context. To this end, we tested the essentiality, specificity and efficacy of CRISPR dependency screens. The top co-dependent genes by CRISPR-Cas9 setting specify the degree of gene essentiality and replication fitness (see Methods). To further inspect the importance of SLC7A11 in colon cancer, we investigated CRISPR-based dependency map for Xc- system (*i.e.,* SLC7A11, SLC3A2) and the CD44 which stabilize Xc- system in the plasma membrane. We compared genes that were characterized by their correlation following CRISPR gene depletion and focused on the 100 top listed correlated genes for each. **Fig. 4A** compared genes that by their overlap among the co-dependent CRISPR screening results. There are 27 genes that are shared by the Xc- system. As expected, the cell-surface glycoprotein CD44 that involves in cells’ interactions, adhesion and migration shows a minimal overlap due to its different functionality. Only SDHA is shared with all three components (including CD44). This gene belongs to the mitochondrial respiratory complex and highlight the importance of mitochondrial function as a central hub for Xc- gene dependency. In contrast, a substantial overlap between the co-dependent SLC7A5 and SLC3A2 sets is observed (63 genes). The detailed results of the Venn analysis are available in Supplementary **Table S5**.

**Figure 4.**
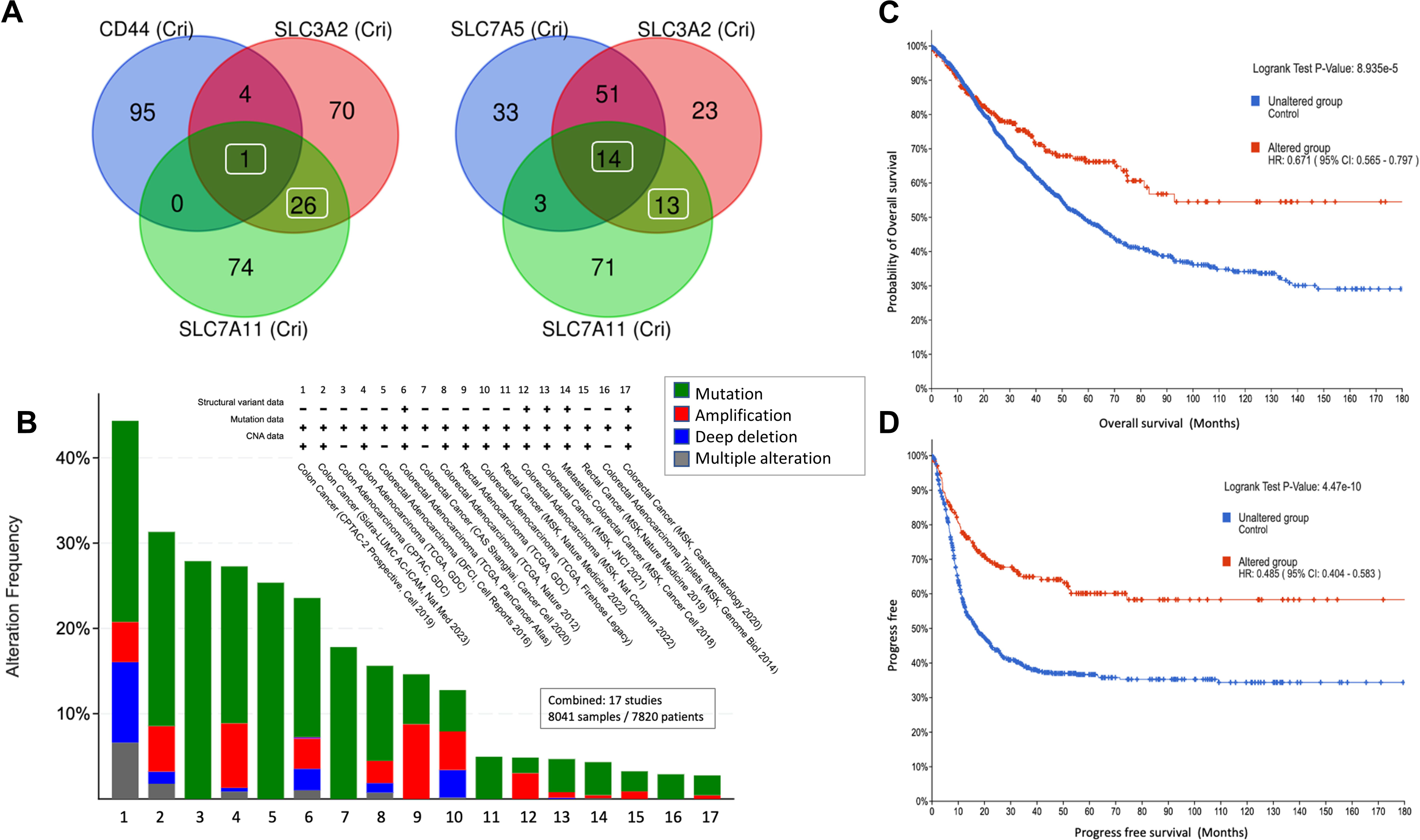
Gene set of CRISPR-induced Xc- fitness and CRC survival. **(A)** Venn diagram for the co-dependent genes by CRISPR-Cas9 setting (DepMap-based, see Methods). Top 100 correlated genes were included in the analysis. There are 27 shared genes (left, white frame) between the two membrane transporters with only 1 overlapped gene also with CD44. In contrast, the SLC7A5 displayed strong shared signal with SLC3A2 (Right, 65 genes) among them 14 genes are shared by all three gene sets. **(B)** Total 17 bowel cancer studies are listed according to the type of alteration in their genes (e.g., mutations, copy number variations; see legend for colors). We selected 17 of 26 studies reported in cBioPortal (a total of 8041 samples). Selected studies have ≥100 samples each (appendiceal cancer cohort was excluded). The survival plots (C and D) indicate the unaltered and altered sets (blue and red, respectively) for samples with alteration in any of the 27 shared genes (shown in A). The statistical significance, Hazard ratio (HR) with 95% confidence is indicated. **(C)** The overall survival plot (OS) for 150 months. **(D)** Progress free survival (PFS) curve for 150 months. Statistically significance is indicated.

For testing the impact of alteration in the overlapping genes (27) on the survival of COAD patients, we created a cohort composed of 17 bowel-derived complementary cancer studies. About 12% of all samples (920 samples) carry alterations in any of the 27 shared gene set (**Fig. 4B**, Supplementary **Table S3**). The overall survival (OS, **Fig. 4C**) and progression-free survival (PFS, **Fig. 4D**) show that for both survival settings, survival of the altered genes is markedly enhanced. The hazard ratio (HR) indicates an improved survival relative to unaltered group (accounts for 88% of the samples). The HR for overall survival (OS) was 0.594 (**Fig. 4C**) and the progression free survival (PFS) was 0.383 (**Fig. 4D**). The results are consistent with the presence of overexpressed level of SLC7A11 in tumors, where alterations in SLC7A11 and its correlated gene set, are linked to improved survival. This is the basis for the importance of SLC7A11 inhibitors in therapy (Bartha & Gyorffy, 2021).

### Functional analysis of gene dependencies in the Xc- system using in silico approach

To further substantiate a functional link between the strongly increased gene expression in COAD and robust results from CRISPR codependency tests (**Table 2**), we seek for genes whose expression is altered with respect to loss of function (LOF) of selected gene set (using MuTarget, see Methods, **Fig. 5**). The logic of this analysis is to provide a more direct connection between damaged genotype and expression of overlooked genes as outcome. Note that the MuTarget approach is indifferent to the expression trend in tumor versus healthy samples. We restricted the analysis to the 5-gene functional set of SLC7A11, SLC3A2, ATIC, TFRC and UMPS (coined FunSet, **Table 2**) only with disrupting damaging mutations. We seek to find genes that were significantly changed in their expression (up- or downregulation) for the genotype-based partition of COAD samples. The results of the six most significant genes are shown in **Fig. 5**. In the collection of samples where SLC7A11 and its associates lost their function due to damaging mutations, SELENBP1 and SGK2 were downregulated while OXCT1 was upregulated. While causality cannot be inferred from such analysis, the affected genes all act in metabolism regulation and epithelial–mesenchymal transition (EMT). Specifically, OXCT1 is a mitochondrial matrix enzyme that plays a central role in ketone catabolism. SELENBP1 is a selenium-binding protein which may be involved in sensing reactive xenobiotics, and SGK2 is a serine/threonine-protein kinase which is involved in the regulation of membrane channels and amino acid transporters. SELENBP1 and SGK2 were associated with their impact of EMT (Liu *et al*, 2020b; Zhang *et al*, 2022) (**Fig. 5A**). The results for MuTarget identified genes are in Supplementary **Table S6**. Whether such gene targets specify the contribution of EMT in CRC calls for further investigation. We applied GO annotation enrichment test whose change in expression was linked to FunSet. The enriched annotations among the upregulated genes support the occurrences of immune response genes. For example, GO_BP, adaptive immune response (p-value 4.4e-34) and antigen processing and presentation of peptide (p-value 9.2e-53). In contrast, the annotations of the downregulated genes (e.g., SGK2) are linked to various cell homeostasis processes including negative regulation of WNT signaling, ion transport, xenobiotic metabolic process and more (see Supplementary **Table S6**). We conclude that via such COAD sample stratification, candidate genes and pathways could lead to drug targeting and molecular manipulation of overlooked targets.

**Figure 5.**
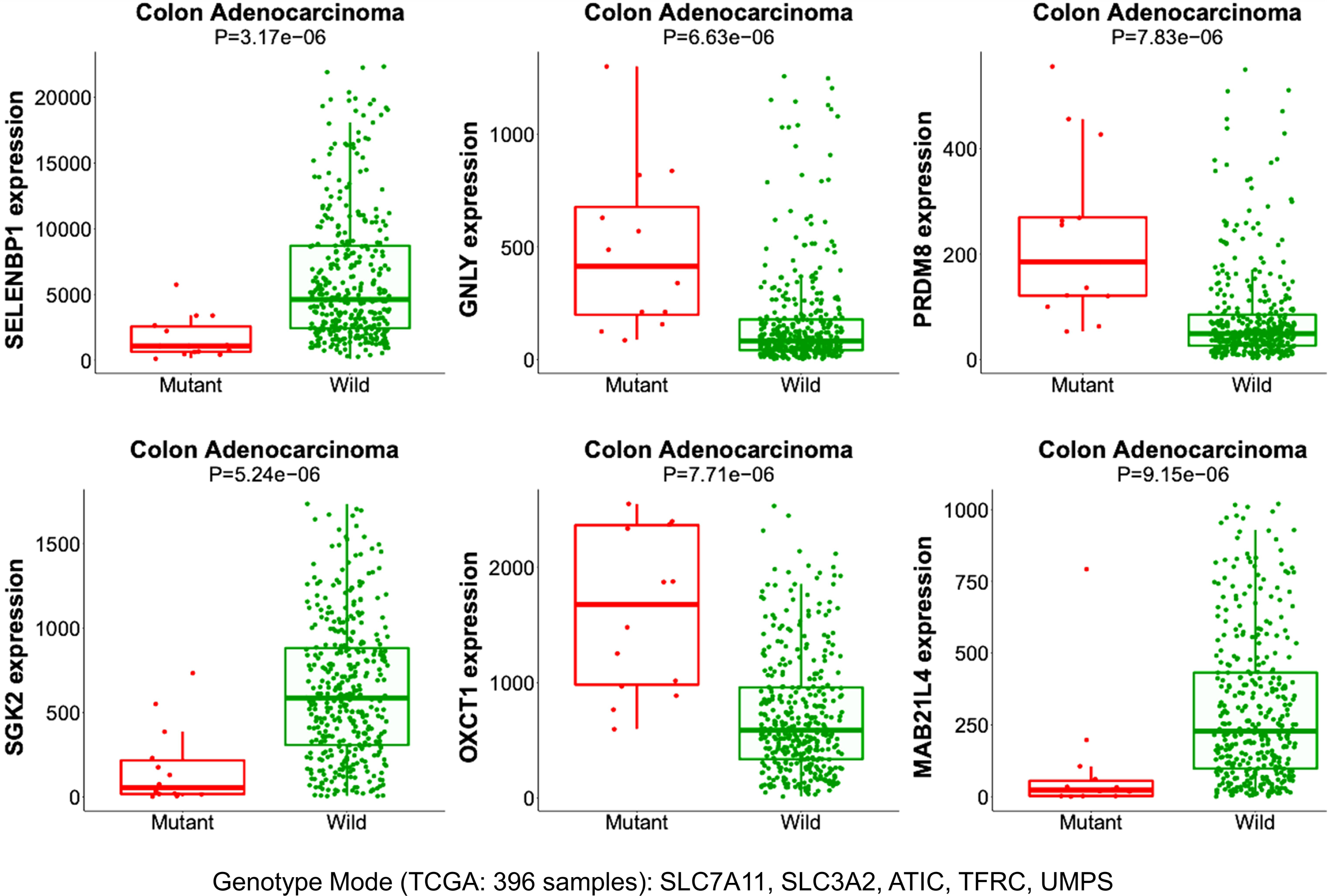
Analysis of MuTarget for paired genomic and transcriptomic data from COAD TCGA samples. Genotype mode. Six most significant altered gene expression related to the combined damaging mutations in SLC7A11, SLC3A2, ATIC, TFRC and UMPS (14 samples). Only significant changes of individual gene expression were considered (FC≥|2|, p-value <1e-3, minimal ≥100 CPM. The reported p-value is based on Mann-Whitney U-test.

**Table 2.** Overlapping CRISPR co-dependent genes of Xc- system genes in COAD samples FC is the fold change. In bold face FC>2.0 for Tumor (T) vs healthy (H). ^b^In bold face genes that amplified the metastatic (Met.) state.

| Gene | Description | Main function | FC<br>T vs H<sup>a</sup> | FC<br>Met. vs<br>T | FC<br>Met. vs<br>H<sup>b</sup> |
| --- | --- | --- | --- | --- | --- |
| *KRT16* | Keratin, type I cytoskeletal 16 | Intermediate filament | **13.13** | 0.13 | 1.73 |
| *SLC7A11* | Cystine/glutamate transporter | Cystine transport as redox regulator | **9.22** | 0.34 | 3.15 |
| *PRH2* | Proline rich protein HaeIII subfamily 2 | Secreted glycoprotein | **3.27** | 0 | 0 |
| *MTHFD1* | Methylenetetrahydrofolate dehydrogenase | De novo purine synthase | **2.72** | 0.62 | 1.7 |
| *ATIC* | Formyltransferase/IMP cyclohydrolase | De novo purine biosynthetic pathway | **2.7** | 2.15 | **5.79** |
| *RPIA* | Ribose 5-phosphate isomerase A | Pentose-phosphate pathway | **2.65** | 0.77 | 2.03 |
| *UMPS* | Uridine monophosphate synthetase | De novo pyrimidine biosynthetic pathway | **2.65** | 6.34 | **16.79** |
| *CAD* | Transcarbamylase-dihydroorotase | De novo biosynthesis of pyrimidine nucleotides | **2.62** | 1 | 2.61 |
| *TFRC* | Transferrin receptor | Cellular iron uptake | **2.51** | 1.91 | **4.79** |
| *RAB36* | Ras related protein Rab-36 | Vesicle-mediated transport | **2.47** | 0.8 | 1.98 |
| *SLC3A2* | 4F2 Cell surface antigen heavy chain | Transport of L-type amino acids | **2.27** | 0.54 | 1.22 |

### Differential protein expression of xCT and its interactors determines chemosensitivity and ferroptosis in PDO models

To validate CRISPR-based functional data, we analyzed newly generated CRC patient-derived organoids (PDOs) to assess the expression of xCT-interacting proteins. This included metabolic profiling, drug response assays targeting xCT inhibition under standard treatment regimens, in PDOs confirmed for genetic fidelity of the PDOs to corresponding tumor counterpart. Clinical parameters of the PDOs used in this study are listed in **Table 3**.

**Table 3.** Clinical parameters of patients with CRC and basic molecular properties of retrieved tissue specimen propagated for *in vitro* model generation

| Tumor code | Primary/<br>metastasis | Gender | Age at surgery | TNM | Grading | MMR | Mut |
| --- | --- | --- | --- | --- | --- | --- | --- |
| 86 | primary | male | 59 | pT4a pN0 L0 V0 Pn0 R1 | G3 | MSS | N.A. |
| 149 | metastasis | female | 78 | pT4b pN2a L1 V1 Pn1 pM1 cR1 | G3 | N.A. | N.A. |
| 151 | primary | male | 74 | pT3 pN1b L1 V0 Pn0 M0 R0 | N.A. | MSS | N.A. |
| 162 | metastasis | male | 54 | pT3 pN2b pM1 L1 V1 Pn1 R1 | G3 | MSS | KRAS |
| 1719 | primary | female | 60 | pT1 pN0 M0 L0 V0 Pn0 R0 | Gx | MSS | N.A. |

Flow cytometric analyses revealed no statistically significant differences in xCT surface expression between tumor and healthy colon organoids, despite slightly elevated mean fluorescence intensity (MFI) values in tumor organoids (**Figs. 6A, 6B**). CRC-derived PDOs demonstrated significantly lower intracellular cystine concentrations and higher levels of unbound Fe²⁺ compared to healthy organoids (p-val=0.018, p-val=0.039, respectively). Although not directly tested in this study, the elevated cystine levels in healthy PDOs may enhance their protection against reactive oxygen species (ROS), while their reduced Fe²⁺ content suggests diminished susceptibility to ferroptosis. Interestingly, no correlation was observed between xCT expression and intracellular cystine levels (r=−0.13, p-val=0.78). However, xCT expression showed a strong and very significantly positive correlation with expression of transferrin receptor (TFRC) levels (r=0.88, p-val=0.009) and unexpectedly, a significant negative correlation with the expression of stem cell marker CD44 (r≈−0.77, p-val=0.042). Additionally, CD44 inversely correlated with intracellular Fe²⁺ levels (r=−0.8, p-val=0.0326), suggesting a potential antagonistic relationship between this marker and iron accumulation **(Fig. 6C).** We concluded that functional analysis using PDOs allows to get a closer view of the tumor pathophysiology and person to person heterogeneity.

**Figure 6.**
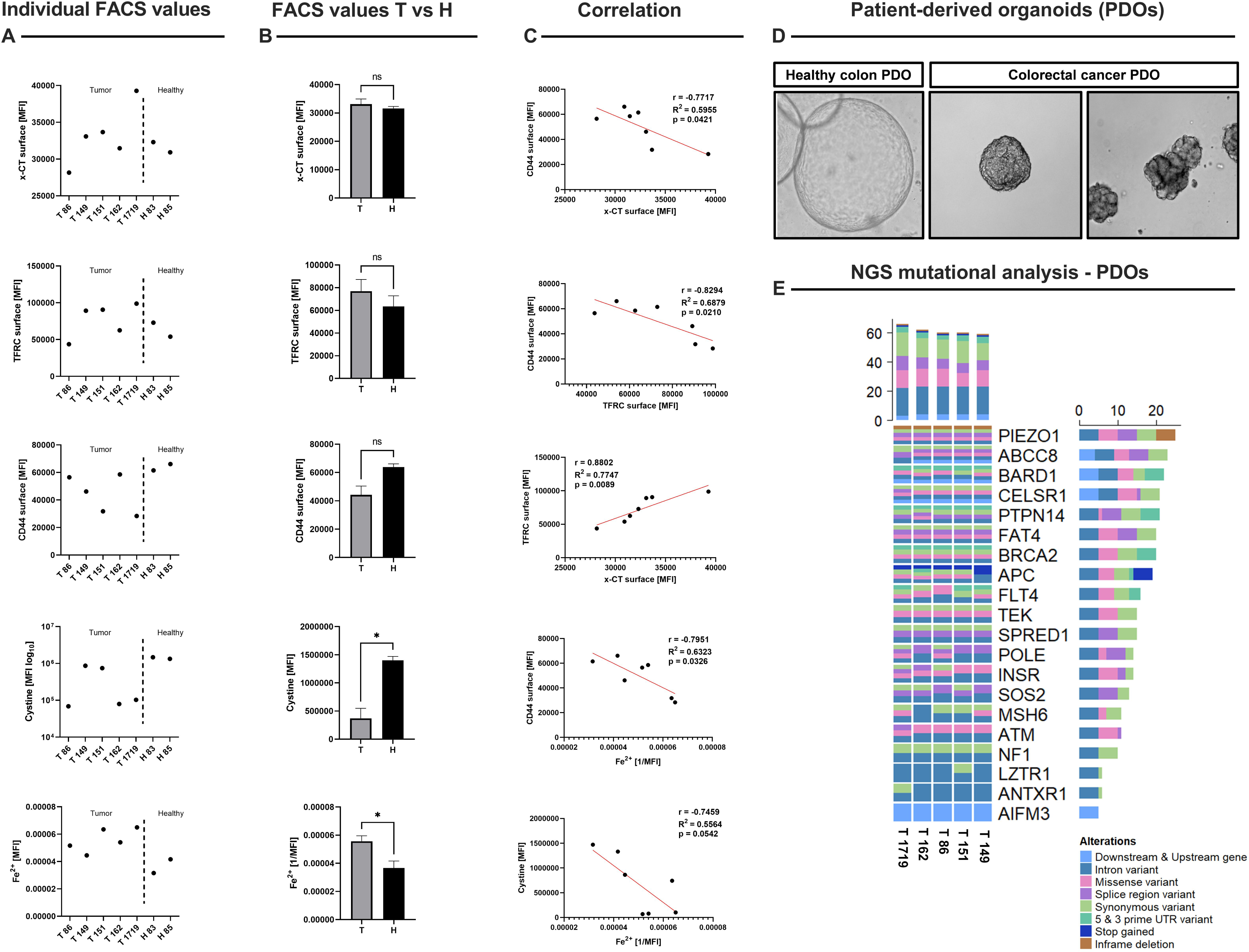
Functional assays using flow cytometric analyses of the protein levels of xCT, TFRC, CD44, and the relative amonuts of cystine, and Fe^2+^ in tumor vs. healthy PDO samples (**A, B**) and their correlations (**C**). T= tumor PDO, H= healthy PDO. (**D**) Exemplary pictures of used PDOs showing different distinct morphologies, with NGS profiles illustrating key mutational landscapes in representative PDOs **(E).** *p-val < 0.05; **p-val < 0.01; ***p-val < 0.001; ****p-val < 0.0001.

Drug viability assays on tumor organoid lines (T86, T162, T149, T1719) revealed differential responses to treatment with standard of care compound 5-fluorouracil (5-FU; 10 µM), Erastin (10 µM), or their combination (**Fig. 7A**). While T86 and T162 exhibited minimal viability reduction (p-val<0.05, p-val>0.05), T149 and T1719 were highly susceptible to all treatments, with partial rescue observed upon co-administration of Erastin (blocks the activity of system Xc⁻) and the iron chelator deferoxamine (DFO) (T149, T162, T1719). Correlative analyses demonstrated that PDOs with elevated xCT surface expression was more sensitive to Erastin (r≈0.95, p-val=0.046) **(Fig. 7B)** and exhibited enhanced sensitivity to 5-FU (r=0.97, p-val=0.034) **(Fig. 7C)**. Elevated TFRC levels were associated with increased sensitivity to Erastin (r≈0.93, p-val≈0.0658), whereas higher CD44 expression correlated negatively with both Erastin efficacy (r≈−0.98, p-val=0.015) and 5-FU-induced cytotoxicity (r≈−0.99, p-val=0.005).

**Figure 7.**
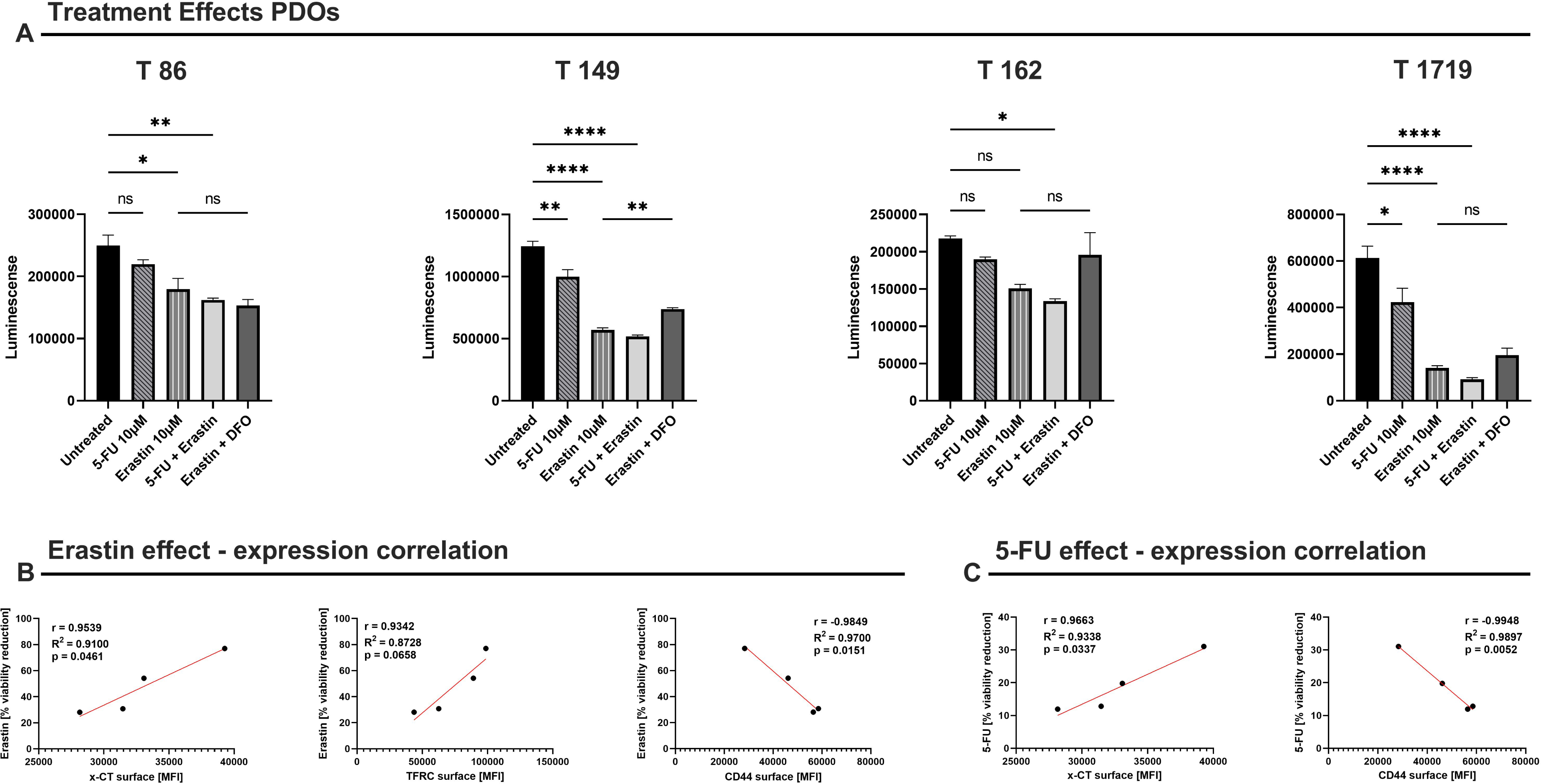
Luminescence-based viability of patient-derived organoids (T86, T149, T162, T1719) after treatment with 5-FU 10 µM and the ferroptosis inducer Erastin 10 µM, either alone or in combination, with or without the iron chelator DFO 100 µM **(A)**. Panels **(B)** and **(C)** show correlations between the baseline expression (MFI) of xCT, TFRC, or CD44 and the percentage viability reduction under Erastin- or 5-FU–based treatments. The p-value thresholds are: *p-val < 0.05; **p-val < 0.01; ***p-val < 0.001; ****p-val < 0.0001.

Collectively, these findings highlight the differential roles of xCT, TFRC, and CD44 in modulating drug responses in CRC PDOs. The marked vulnerability of T149 and T1719 organoids to Erastin- and 5-FU-induced cell death underscores the therapeutic potential of targeting these pathways.

### Direct effect of differentially expressed miR-135b in CRC samples

Insights from post-transcriptional regulation can bridge Xc- system overexpression with tumorigenesis. An attractive mechanism concerns the epithelial to mesenchymal transition (EMT), where specific TFs are regulated through cellular miRNAs along cancer progression (Vu & Datta, 2017). The mode of action of miRNAs involves direct and indirect effects. The direct effect is by targeting mRNA through pairing with the 3’-UTR miRNA binding sites (MBS), while indirect effects are manifested through competition on MBS (i.e., ceRNA) and via TF regulation that drives global changes in chromatin structure and cell decisions. To provide a further support for the regulation layers in CRC samples with respect to the Xc- system, we determined the miRNA profiles for each of the reported 32 patients by comparing healthy and tumor samples in pairs. Among the 450 identified miRNAs (Supplementary **Table S7**), there were 40 downregulated and 20 upregulated miRNAs (p-val FDR: <5e-05; fold >|2.4|; coined differentially expressed miRNAs, DEMs). As in many tissues, only a handful of miRNAs accounts for the majority of miRNA (about 20 miRNAs accounts for 80% of miRNA expression). From this list of top expression miRNAs, hsa-miR-21-5p (accounts for 7.1%) was upregulated among DEMs (Supplementary **Table S7**).

**Table 3** lists 10 most significant DEMs for up- and downregulated miRNAs (5 miRNAs each) along with their abundance. In most cases (80%), the expression trend of our CRC cohort is in agreement with the meta-analysis validated results (based on miRTarBase (Huang *et al*, 2022)). The most significant upregulated miRNA is miR-135-5p (19.1 folds) which is associated with a ranked list of targets (804, by miRDB) (Chen & Wang, 2020). By focusing on the top 5% of ranked targets, we identified a few genes which were downregulated in our CRC samples (downregulated DEGs, Supplementary **Table S2**). These target candidates for miR-135b-5p were NR3C2, SLC30A4, and KLF4. NR3C2 and KLF4 were strongly downregulated (4.2 and 3.6 folds respectively) in our CRC cohort.

The other miRNAs that are strongly upregulated are miR-96-5p and miR-1246 (**Table 4**). Multiple studies confirm that miR-96-5p is overexpressed in CRC and enhances tumor cell viability, colony formation, and cell cycle progression. Our computational predictions provide a possible gene-TF-miRNA network that indicate SLC7A11 with transcription factors involved in core biological processes such as ZEB1, TCF3 and LEF1. The gene-TF-miRNA network is available in Supplementary **Fig. S2.**

**Table 4.** Upregulation and downregulation miRNAs from 32 CRC patients

| miRNA | Precursor | Trend | Amount | FDR | Fold<br>(T vs H) | Meta<sup>a</sup><br>(TarBase) |
| --- | --- | --- | --- | --- | --- | --- |
| miR-1-3p | mir-1-1; mir-1-1 | Down | 493.2 | 7.2E-12 | 0.20 | Down |
| miR-1246 | mir-1246 | Up | 48.4 | 8.9E-11 | 6.78 | Up |
| miR-129-5p | mir-129-1; mir-129-2 | Down | 112.7 | 1.5E-14 | 0.12 | Up |
| miR-133a-3p | mir-133a-1; mir-133a-2 | Down | 59.5 | 7.3E-18 | 0.12 | Down |
| miR-135b-5p | mir-135b | Up | 185.2 | 3.1E-29 | 19.12 | Up |
| miR-139-5p | mir-139 | Down | 397.4 | 4.7E-21 | 0.20 | Down |
| miR-183-5p | mir-183 | Up | 1372.4 | 1.1E-08 | 3.82 | Up |
| miR-215-3p | mir-215 | Down | 43.4 | 1.6E-08 | 0.13 | Down |
| miR-7974 | mir-7974 | Up | 20.9 | 1.1E-07 | 5.55 | Same |
| miR-96-5p | mir-96 | Up | 205.0 | 1.6E-08 | 4.26 | Up |

### Differentially expressed miRNAs support mitochondrial and mitotic signals in colon

Most effects of miRNAs are mediated indirectly. Specifically, we inspected the potential indirect effect of miRNAs for the set of DEMs from our CRC cohort, by an unbiased annotation enrichment test. **Fig. 8A** shows results of a meta-analysis of miRNA expression according to a large compilation of GEO experiments comparing tumor to healthy samples across various neoplasms. The miRNAs are sorted according to the fold change reported in our cohort (Supplementary **Table S7**). We show a substantial agreement between miRNA ranking in Magdeburg patient data to the public COAD dataset. However, the assignment of the miRNA for ranked up- and downregulation for other cancer types (liver, pancreas, thyroid) is far less consistent. **Fig. 8A** demonstrates that each cancer type is specified by its unique miRNA expression profile (Mahlab-Aviv *et al*, 2021).

**Figure 8.**
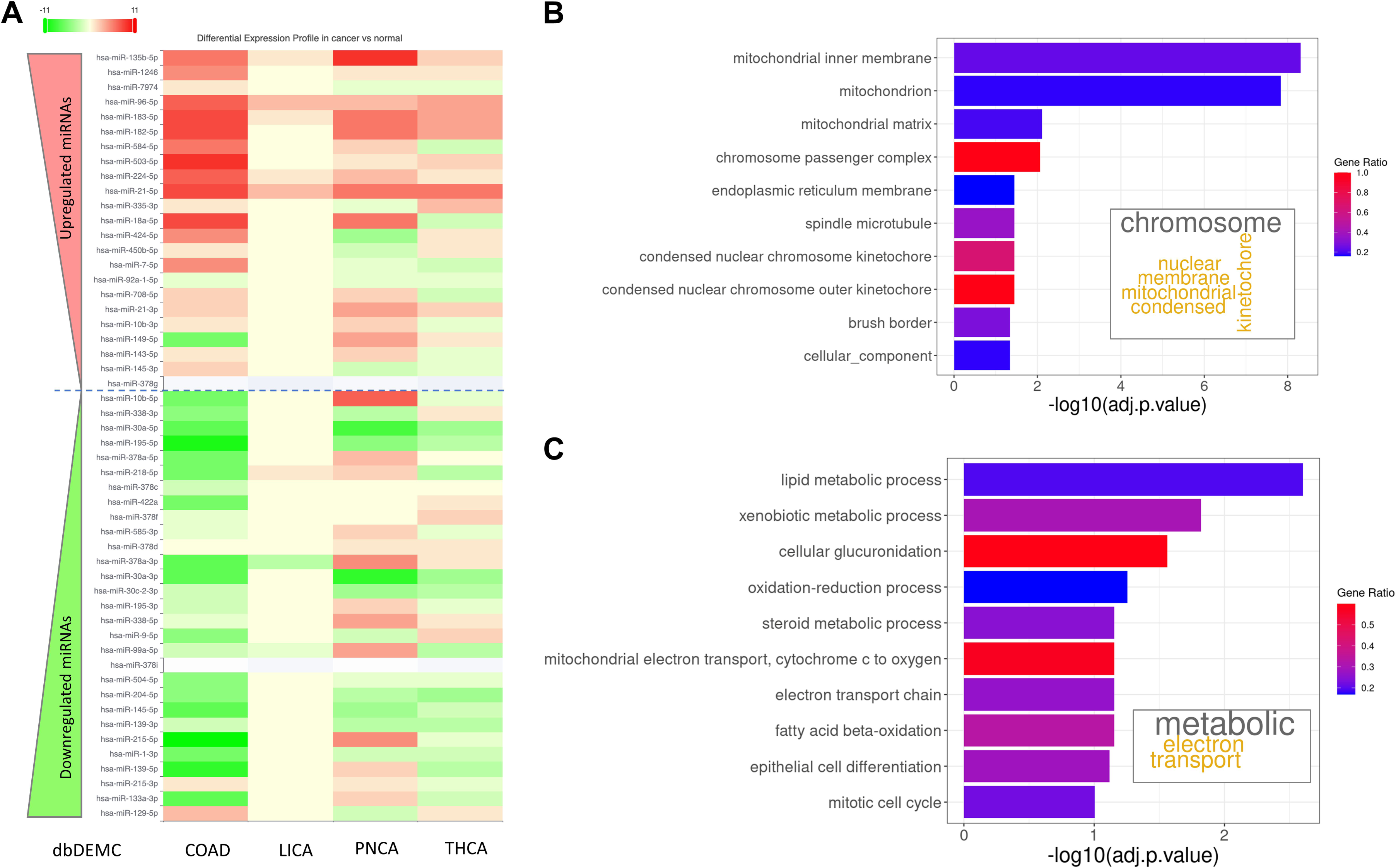
DEM profile and Functional enrichment**. (A)** Results of meta-analysis of miRNAs in 4 different cancer types (colon adenocarcinoma, COAD; cancer of liver cancer (LICA), pancreas (PNCA) and thyroid (THCA). The rank of DEM (total 52 different miRNAs) is according to the fold change from up- to downregulated miRNAs. The meta-analysis was performed by expression miRNA compilation by dbDEMC (see Methods). The dashed horizontal line separates up and downregulated miRNA in from the CRC cohorts (Supplemental **Table S7**). Enrichment scheme for GO annotation by miRtoGO (see Methods) with 10 top DEM (from Table 4). **(B)** Enrichment scheme for GO_CC. **(C)** Enrichment scheme for GO_BP. The histogram is sorted by the significance by -log10 (adjusted p-value) for each of the enriched term. The color indicates the fraction of genes included in each term from the total number of genes associated with the indicated term. Inset, a word map of the 10 most significant terms.

The bar plots show the statistical enrichment for GO annotations by indirect effect of miRNAs in colon tissue (**Fig. 8B, 8C**). The input for the enrichment is the 10 top DEMs (**Table 4**). **Fig. 8B** shows an enrichment plot for GO cellular components (GO_CC) which indicates a strong signal for mitochondria, membrane and mitotic components. **Fig. 8C** shows the enrichment of GO biological processes (GO_BP) that support metabolic alteration and oxidative processes but also cell cycle. We propose that the observed set of miRNAs expose indirect regulation that may drive cells to adapt an alternative metabolic state for electron transfer and oxidation regulation.

## Discussion

Aberrant activation of SLC7A11 is involved in various cancer types, where it influences redox balance, metabolic flexibility, immune function, and ferroptosis (Jyotsana *et al*., 2022). We have focused on the role of the Xc- system in CRC cancer and showed that it acts as a hub connecting an elaborate signature of mitotic cells and the cell cycle with a metabolic program (Fig. 2B). SLC7A11 may have conflicting effects on different cancer cells. For example, in glioblastoma, the overexpression of the Xc- system induces oxidative stress and apoptosis, in contrast to CRC, where it is a strong mediator for cell viability. Clinically, suppression of SLC7A11 function in CRC (*e.g.,* by p53 or BECN1) can activate ferroptosis, which makes the tumor sensitive to radiotherapy. To further analyze the cellular role of Xc-, we inspected the downstream glutathione pathways (e.g., GPX4, GPX8). We observed that the expression levels of GPX genes remained stable within the cohort of 32 patients (Supplementary Table S1). Essential and co-essential genes across many cell lines are a useful approach to identifying shared functional pathways (Arnold *et al*, 2022). The CRISPR-based platform for GPX4 and GPX8 across 39 cells, which originated from COAD, did not support such a coherence signature of co-dependency. These results argue for the involvement of alternative pathways for Xc- system driven tumorigenesis. In contrast, SLC7A5 was exceptionally correlated with SLC3A2 (Fig. 4A). The role of SLC7A5 in CRC is to maintain intracellular amino acid levels following KRAS activation through transcriptional and metabolic reprogramming (Najumudeen *et al*, 2021).

Among the genes that strongly correlated with the Xc- system in CRISPR-based assays, TFRC (transferrin receptor) was found to be upregulated in COAD and implicated in promoting metastatic tumors (Table 2). Due to its role in iron accumulation and activation, TFRC also promotes nucleotide biosynthesis, DNA repair, and cell survival in colorectal tumors (Schwartz *et al*, 2021). Additionally, impairing TFRC function has been suggested as a target for inhibiting tumor growth. Specifically, reducing the iron influx can induce DNA damage and apoptosis (Kim *et al*, 2023). The interplay between TFRC and SLC7A11 provides cancer cells with a survival advantage based on these findings. We suggest that altering iron homeostasis and redox balance through manipulating the levels of TFRC and SLC7A11 can serve as potential therapeutic targets.

To further elucidate the functional role of SLC7A11 in COAD we employed a short-term cultured PDO platform. Quality control of generated PDOs reveals typical COAD specific mutation profiles (**Figs. 6D, 6E**) and allows comparison of in vitro to clinical drug responses in dependency of relevant driver mutations. Recent research by De et al. (De *et al*, 2025) highlights the potential of targeting patient-specific KRAS G12D mutations to promote ferroptosis, particularly when combined with other ferroptosis inducers. This approach strengthens the clinical relevance of xCT as a therapeutic target and underscores the importance of tailoring treatment strategies to individual mutation profiles. Our results add new knowledge proposing a particular centrality of xCT activity in CRC by demonstrating how differential xCT cell surface expression modulates metabolic pathways and therapeutic. Consistent with previous studies showing that xCT regulates redox balance and ferroptosis in various malignancies, our data reveal a paradoxical finding. The CRC organoids displaying higher xCT expression exhibited enhanced sensitivity to 5-FU, contradicting the conventional view that xCT upregulation confers chemotherapy resistance. A plausible explanation for this discrepancy may lie in the heightened metabolic demands and redox flux imposed by xCT overexpression. Under certain conditions, this pro-survival mechanism can render cells more vulnerable to genotoxic or ferroptosis-inducing stress. Notably, similar observations in glioblastoma demonstrate that high xCT levels elevate intracellular oxidative stress and trigger apoptosis, especially under combined stressors such as chemotherapy or glutathione depletion (Liu *et al*, 2020a). In ovarian cancer, elevated SLC7A11 was associated with improved sensitivity to paclitaxel therapy and superior OS, potentially due to increased autophagy under chemotherapeutic stress via competing endogenous RNA mode of action (ceRNA) (Ke *et al*, 2021). Taken together, these findings emphasize that while xCT overexpression often supports tumor survival, it can also sensitize cells to cytotoxic agents if oxidative or replicative stress surpasses the cellular compensatory capacity. In addition, work by Gu et al. linked metabolic burden and proliferation, it was shown that mTORC2, a critical regulator of amino acid metabolism can phosphorylate xCT and therefore, inhibiting its activity (Gu *et al*, 2017). From a technical perspective, we acknowledge that our selected method of antibody-based quantitation of xCT in living cell models was restricted to cell surface epitope exposure. Notably, cell surface localization of xCT is subjected to regulation and to the cell metabolic state (Koppula *et al*., 2021). While using the PDO platform to validate our findings enables a more personalized approach with focus on stem cell population, analyzing bulk RNA sequencing data often ignores the biological highly relevant intra-tumoral heterogeneity. This might explain why our analysis in public dataset could not identify a statistically significant correlation between patientś response to chemotherapy and x-CT transcript abundancy (**Fig. S2**). Nevertheless, our data underscore the tumor- and context-specific roles of xCT in chemotherapeutic responses.

Increased TFRC promotes cellular iron uptake, which can support DNA synthesis and repair but may also facilitate ferroptosis under conditions of redox dysregulation. We identified that cells with high xCT expression possess also TFRC protein upregulation, further suggesting a convergence between iron metabolism and the xCT system. Consequently, manipulating iron homeostasis in tandem with xCT and combined standard chemotherapy treatments appear to be a promising therapeutic strategy (Lei *et al*, 2022). While xCT has been shown to be essential in multiple contexts for sustaining tumor growth (Muir & Vander Heiden, 2018), our finding that 5-FU retained (and even enhanced) efficacy in xCT-high organoids illustrates how metabolic pathways can exert dual-edged effects in CRC. These observations align with emerging data from other tumor types, indicating that SLC7A11 can either promote or hinder survival depending on specific cellular conditions (Chen *et al*, 2009). By synthesizing transcriptomic, CRISPR-based, metabolic and functional cell model evidence, it becomes clear that post-transcriptional and post-translational modifications, as well as more comprehensive metabolomic analyses, are essential to fully decipher how SLC7A11 and its partner molecules govern CRC progression and response to therapy.

In summary, we propose that CRC patients prone to receive neoadjuvant chemotherapy may benefit from pre-treatment screenings to quantify xCT activation and subsequent stratification into high and low responsiveness to standard chemo therapy treatment, which may lead to individually adjusted therapy regimes. We highlight the possibility to improve standard of care chemotherapy when combining therapy with xCT inhibitor treatment in tumors with high xCT activation. Moreover, our epigenetic analysis reveals hitherto under-investigated microRNA targets. Studying their role in CRC may help to improve future colon cancer care, by epigenetic interference in colon cancer oxidative stress homeostasis. Further investigations, especially using functional assays on pathophysiological predictive model systems and genetic modeling, are warranted to probe for any biological relevance our hypothesis.

## Ethics approval and consent to participate

The Ethics Committee of University Magdeburg approved this study under registry 33/01 and 46/22.

## Abbreviations

CD44v: CD44 variant
COAD: colorectal adenocarcinoma
CPM: counts per million
CRC: colorectal cancer
DEG: differential expression gene
DEM: differential expression miRNA
EMT.: epithelial–mesenchymal transition
FC: fold change
GO: gene ontology
GPX: glutathione peroxidases
GSH: glutathione
GTEx: genotype tissue expression
ICI: immune checkpoint inhibitor
LOF: loss of function
MFI: mean fluorescence intensity
NGS: Next generation sequencing
OS: overall survival
PCA: principal component analysis
PDO: patient-derived organoid
PFS: progression-free survival
READ: rectum adenocarcinoma
RNA-seq: RNA sequencing
ROS: reactive oxygen species
TCGA: the cancer genome atlas
TMM: trimmed mean of the M-values
TPM: transcript per million
xCT: solute carrier family 7, member 11

## References

Arnold PK, Jackson BT, Paras KI, Brunner JS, Hart ML, Newsom OJ, Alibeckoff SP, Endress J, Drill E, Sullivan LB et al (2022) A non-canonical tricarboxylic acid cycle underlies cellular identity. Nature 603: 477–481

Bartha A, Gyorffy B (2021) TNMplot.com: A Web Tool for the Comparison of Gene Expression in Normal, Tumor and Metastatic Tissues. Int J Mol Sci 22

Biller LH, Schrag D (2021) Diagnosis and treatment of metastatic colorectal cancer: a review. Jama 325: 669–685

Bonifácio VD, Pereira SA, Serpa J, Vicente JB (2021) Cysteine metabolic circuitries: Druggable targets in cancer. British journal of cancer 124: 862–879

Chen R, Song Y, Zhou Z, Tong T, Li Y, Fu M, Guo X, Dong L, He X, Qiao H (2009) Disruption of xCT inhibits cancer cell metastasis via the caveolin-1/β-catenin pathway. Oncogene 28: 599–609

Chen Y, Wang X (2020) miRDB: an online database for prediction of functional microRNA targets. Nucleic acids research 48: D127–D131

Cheng X, Wang Y, Liu L, Lv C, Liu C, Xu J (2022) SLC7A11, a potential therapeutic target through induced ferroptosis in colon adenocarcinoma. Frontiers in Molecular Biosciences 9: 889688

Choi A, Jang I, Han H, Kim MS, Choi J, Lee J, Cho SY, Jun Y, Lee C, Kim J et al (2021) iCSDB: an integrated database of CRISPR screens. Nucleic Acids Res 49: D956–D961

Consortium G (2013) The Genotype-Tissue Expression (GTEx) project. Nat Genet 45: 580–585

Daher B, Vučetić M, Pouysségur J (2020) Cysteine depletion, a key action to challenge cancer cells to ferroptotic cell death. Frontiers in Oncology 10: 723

De D, Cohen D, Mitani Y, Li T, Bethancourt C-N, Gabre J, 2025. Sensitizing K-Ras G12D mutant colorectal patient-derived organoids via ferroptosis pathway. American Society of Clinical Oncology.

Del Toro N, Shrivastava A, Ragueneau E, Meldal B, Combe C, Barrera E, Perfetto L, How K, Ratan P, Shirodkar G (2022) The IntAct database: efficient access to fine-grained molecular interaction data. Nucleic acids research 50: D648–D653

Dempster JM, Rossen J, Kazachkova M, Pan J, Kugener G, Root DE, Tsherniak A (2019) Extracting biological insights from the project achilles genome-scale CRISPR screens in cancer cell lines. BioRxiv: 720243

Fekete JT, Gyorffy B (2023) New Transcriptomic Biomarkers of 5-Fluorouracil Resistance. Int J Mol Sci 24

Fekete JT, Győrffy B (2019) ROCplot. org: Validating predictive biomarkers of chemotherapy/hormonal therapy/anti-HER2 therapy using transcriptomic data of 3,104 breast cancer patients. International journal of cancer 145: 3140–3151

Gu Y, Albuquerque CP, Braas D, Zhang W, Villa GR, Bi J, Ikegami S, Masui K, Gini B, Yang H et al (2017) mTORC2 Regulates Amino Acid Metabolism in Cancer by Phosphorylation of the Cystine-Glutamate Antiporter xCT. Mol Cell 67: 128–138 e127

Hanahan D (2022) Hallmarks of cancer: new dimensions. Cancer discovery 12: 31–46

He J, Ding H, Li H, Pan Z, Chen Q (2021) Intra-tumoral expression of SLC7A11 is associated with immune microenvironment, drug resistance, and prognosis in cancers: a pan-cancer analysis. Frontiers in Genetics 12: 770857

Huang H-Y, Lin Y-C-D, Cui S, Huang Y, Tang Y, Xu J, Bao J, Li Y, Wen J, Zuo H (2022) miRTarBase update 2022: an informative resource for experimentally validated miRNA–target interactions. Nucleic acids research 50: D222–D230

Jyotsana N, Ta KT, DelGiorno KE (2022) The Role of Cystine/Glutamate Antiporter SLC7A11/xCT in the Pathophysiology of Cancer. Front Oncol 12: 858462

Ke Y, Chen X, Su Y, Chen C, Lei S, Xia L, Wei D, Zhang H, Dong C, Liu X (2021) Low expression of SLC7A11 confers drug resistance and worse survival in ovarian cancer via inhibition of cell autophagy as a competing endogenous RNA. Frontiers in Oncology 11: 744940

Kim H, Villareal LB, Liu Z, Haneef M, Falcon DM, Martin DR, Lee HJ, Dame MK, Attili D, Chen Y et al (2023) Transferrin Receptor-Mediated Iron Uptake Promotes Colon Tumorigenesis. Adv Sci (Weinh) 10: e2207693

Koppula P, Zhuang L, Gan B (2021) Cystine transporter SLC7A11/xCT in cancer: ferroptosis, nutrient dependency, and cancer therapy. Protein & cell 12: 599–620

Lavoro A, Falzone L, Tomasello B, Conti GN, Libra M, Candido S (2023) In silico analysis of the solute carrier (SLC) family in cancer indicates a link among DNA methylation, metabolic adaptation, drug response, and immune reactivity. Front Pharmacol 14: 1191262

Lei G, Zhuang L, Gan B (2022) Targeting ferroptosis as a vulnerability in cancer. Nature Reviews Cancer 22: 381–396

Li T, Fu J, Zeng Z, Cohen D, Li J, Chen Q, Li B, Liu XS (2020) TIMER2. 0 for analysis of tumor-infiltrating immune cells. Nucleic acids research 48: W509–W514

Li Y, Li Y, Xia Z, Zhang D, Chen X, Wang X, Liao J, Yi W, Chen J (2021) Identification of a novel immune signature for optimizing prognosis and treatment prediction in colorectal cancer. Aging (Albany NY) 13: 25518

Liberzon A, Birger C, Thorvaldsdóttir H, Ghandi M, Mesirov JP, Tamayo P (2015) The molecular signatures database hallmark gene set collection. Cell systems 1: 417–425

Lin W, Wang C, Liu G, Bi C, Wang X, Zhou Q, Jin H (2020) SLC7A11/xCT in cancer: biological functions and therapeutic implications. American journal of cancer research 10: 3106

Liu J, Xia X, Huang P (2020a) xCT: a critical molecule that links cancer metabolism to redox signaling. Molecular Therapy 28: 2358–2366

Liu Y, Liu J, Kong X, Li H, Shao J, Jiang Z (2020b) SGK2 is overexpressed in colon cancer and promotes epithelial-mesenchymal transition in colon cancer cells. Eur J Surg Oncol 46: 1912–1917

Lu P, Yu Z, Wang K, Zhai Y, Chen B, Liu M, Xu P, Li F, Zhao Q (2022) DDX21 Interacts with WDR5 to Promote Colorectal Cancer Cell Proliferation by Activating CDK1 Expression. J Cancer 13: 1530–1539

Mahlab-Aviv S, Linial N, Linial M (2021) miRNA Combinatorics and its Role in Cell State Control—A Probabilistic Approach. Frontiers in Molecular Biosciences 8: 772852

Menyhárt O, Győrffy B (2021) Multi-omics approaches in cancer research with applications in tumor subtyping, prognosis, and diagnosis. Computational and structural biotechnology journal 19: 949–960

Miao Y, Ha A, de Lau W, Yuki K, Santos AJM, You C, Geurts MH, Puschhof J, Pleguezuelos-Manzano C, Peng WC et al (2020) Next-Generation Surrogate Wnts Support Organoid Growth and Deconvolute Frizzled Pleiotropy In Vivo. Cell Stem Cell 27: 840–851 e846

Muir A, Vander Heiden MG (2018) The nutrient environment affects therapy. Science 360: 962–963

Nagy A, Gyorffy B (2021) muTarget: A platform linking gene expression changes and mutation status in solid tumors. Int J Cancer 148: 502–511

Najumudeen AK, Ceteci F, Fey SK, Hamm G, Steven RT, Hall H, Nikula CJ, Dexter A, Murta T, Race AM et al (2021) The amino acid transporter SLC7A5 is required for efficient growth of KRAS-mutant colorectal cancer. Nat Genet 53: 16–26

Narasimhan V, Wright JA, Churchill M, Wang T, Rosati R, Lannagan TR, Vrbanac L, Richardson AB, Kobayashi H, Price T (2020) Medium-throughput drug screening of patient-derived organoids from colorectal peritoneal metastases to direct personalized therapy. Clinical Cancer Research 26: 3662–3670

Parker JL, Deme JC, Kolokouris D, Kuteyi G, Biggin PC, Lea SM, Newstead S (2021) Molecular basis for redox control by the human cystine/glutamate antiporter system xc(). Nat Commun 12: 7147

Sayed M, Park JW (2023) miRinGO: Prediction of biological processes indirectly targeted by human microRNAs. Non-coding RNA 9: 11

Schwartz AJ, Goyert JW, Solanki S, Kerk SA, Chen B, Castillo C, Hsu PP, Do BT, Singhal R, Dame MK et al (2021) Hepcidin sequesters iron to sustain nucleotide metabolism and mitochondrial function in colorectal cancer epithelial cells. Nat Metab 3: 969–982

Sugano K, Maeda K, Ohtani H, Nagahara H, Shibutani M, Hirakawa K (2015) Expression of xCT as a predictor of disease recurrence in patients with colorectal cancer. Anticancer Research 35: 677–682

Szklarczyk D, Gable AL, Nastou KC, Lyon D, Kirsch R, Pyysalo S, Doncheva NT, Legeay M, Fang T, Bork P (2021) The STRING database in 2021: customizable protein–protein networks, and functional characterization of user-uploaded gene/measurement sets. Nucleic acids research 49: D605–D612

Tang Z, Kang B, Li C, Chen T, Zhang Z (2019) GEPIA2: an enhanced web server for large-scale expression profiling and interactive analysis. Nucleic Acids Res 47: W556–W560

Testa U, Pelosi E, Castelli G (2018) Colorectal cancer: genetic abnormalities, tumor progression, tumor heterogeneity, clonal evolution and tumor-initiating cells. Medical Sciences 6: 31

Thul PJ, Lindskog C (2018) The human protein atlas: a spatial map of the human proteome. Protein Science 27: 233–244

Vega P, Valentin F, Cubiella J (2015) Colorectal cancer diagnosis: Pitfalls and opportunities. World journal of gastrointestinal oncology 7: 422

Vu T, Datta PK (2017) Regulation of EMT in Colorectal Cancer: A Culprit in Metastasis. Cancers (Basel) 9

Vučetić M, Cormerais Y, Parks SK, Pouysségur J (2017) The central role of amino acids in cancer redox homeostasis: vulnerability points of the cancer redox code. Frontiers in oncology 7: 319

Xu F, Wang Y, Ling Y, Zhou C, Wang H, Teschendorff AE, Zhao Y, Zhao H, He Y, Zhang G et al (2022) dbDEMC 3.0: Functional Exploration of Differentially Expressed miRNAs in Cancers of Human and Model Organisms. Genomics Proteomics Bioinformatics 20: 446–454

Zhang X, Hong R, Bei L, Hu Z, Yang X, Song T, Chen L, Meng H, Niu G, Ke C (2022) SELENBP1 inhibits progression of colorectal cancer by suppressing epithelial-mesenchymal transition. Open Med (Wars) 17: 1390–1404

Zhao Z, Zhou W, Han Y, Peng F, Wang R, Yu R, Wang C, Liang H, Guo Z, Gu Y (2017) EMT-Regulome: a database for EMT-related regulatory interactions, motifs and network. Cell death & disease 8: e2872–e2872

Zygulska AL, Pierzchalski P (2022) Novel Diagnostic Biomarkers in Colorectal Cancer. Int J Mol Sci 23

